# Osteopontin Mediates Uterine Artery Dysfunction in Hypertensive Pregnancy

**DOI:** 10.64898/2026.06.02.729716

**Authors:** Megan L. Lave, Jordan Jones, Violet S. Patterson, Stella Z. Li, Martin W. McBride, Delyth Graham, James C. Lacefield, Genevieve E. Eastabrook, Stephen J. Renaud

**Affiliations:** Department of Anatomy and Cell Biology, Schulich School of Medicine and Dentistry, Western University, London, Ontario, Canada; School of Cardiovascular and Metabolic Health, University of Glasgow, Glasgow, United Kingdom; Department of Electrical and Computer Engineering and School of Biomedical Engineering, Faculty of Engineering, Western University, London, Ontario, Canada; Department of Medical Biophysics and Robarts Research Institute, Schulich School of Medicine and Dentistry, Western University, London, Ontario, Canada; Department of Obstetrics and Gynaecology, Schulich School of Medicine and Dentistry, Western University, London, Ontario, Canada; Children’s Health Research Institute, London Health Sciences Centre Research Institute, London, Ontario, Canada

## Abstract

**Background:** Successful pregnancy requires substantial maternal cardiovascular adaptation, including expansion and remodelling of the uterine arteries to support increased nutrient flow to the fetus. Hypertension is associated with impaired uterine artery remodelling and increased risk of fetal growth restriction and other adverse outcomes, yet the mechanisms driving vascular dysfunction in hypertensive pregnancy remain incompletely understood. Osteopontin, a matricellular protein, is implicated in vascular pathology in hypertension, positioning it as a candidate mediator of impaired uterine artery adaptation during hypertensive pregnancy.

**Methods:** Uterine artery remodelling was compared between pregnant normotensive Wistar-Kyoto (WKY) rats and spontaneously hypertensive stroke-prone rats (SHRSP), a model of chronic hypertension. To determine the role of osteopontin in this process, an osteopontin-deficient SHRSP strain was characterized. Uterine artery blood flow was assessed by Doppler ultrasound, arterial structure was evaluated by histology, and molecular differences were identified by RNA sequencing. Fetal weight and length were measured at mid and late gestation.

**Results:** Compared with WKY, SHRSP fetuses were smaller, and uterine arteries exhibited inward hypertrophic remodelling, characterized by increased wall thickness, reduced lumen area, and elevated resistance index. SHRSP uterine arteries also showed increased expression of inflammatory and vascular pathology-associated genes, including osteopontin. In osteopontin-deficient SHRSP, uterine arteries had larger lumen areas, decreased resistance index, and reduced expression of inflammation-associated genes. Fetal growth was also improved in osteopontin-deficient SHRSP pregnancies.

**Conclusions:** These findings identify osteopontin as a contributor to impaired uterine artery adaptation and suggest that reduced osteopontin may improve vascular remodelling and fetal growth in hypertensive pregnancies.

## Introduction

Pregnancy has been described as the ultimate physiological stress test for the maternal cardiovascular system. To support fetal development, maternal hemodynamics and vascular structure undergo extensive remodelling to accommodate increased blood volume and ensure adequate uteroplacental perfusion (1). Failure to achieve these adaptations can result in serious maternal and fetal complications (2). Hypertensive disorders of pregnancy, which affect approximately 18 million pregnancies globally each year (3), are the most common medical complication of pregnancy and a leading cause of maternal and perinatal morbidity and mortality (4,5). These disorders, which include chronic hypertension, gestational hypertension, preeclampsia/eclampsia, and superimposed preeclampsia, are strongly associated with preterm birth and fetal growth restriction, increasing the risk of neonatal mortality (6,7). Moreover, hypertension during pregnancy confers long-term cardiovascular risk to both mother and child, linking pregnancy complications to future disease (8). This clinical burden is increasing, driven in part by rising rates of cardiometabolic disease in women of childbearing age (9–11). Despite its prevalence and long-term consequences, how hypertension impairs maternal cardiovascular adaptation during pregnancy remains unclear.

A key requirement for successful pregnancy is the remodelling of the uterine arteries, which must expand considerably to accommodate high-volume blood flow to the uterus and placenta (4,12). This transformation is achieved through outward hypertrophic remodelling, characterized by expansion of vessel and lumen cross-sectional area with minimal change in wall thickness (12,13). In hypertensive pregnancies, this process is frequently impaired, as evidenced by elevated resistance indices and diastolic notching on Doppler ultrasound, reflecting compromised uteroplacental perfusion (14). Such vascular dysfunction limits blood flow to the uterus and placenta, and is strongly associated with adverse outcomes, including superimposed preeclampsia and fetal growth restriction (15). However, the mechanisms driving uterine artery dysfunction in hypertensive pregnancies remain poorly defined.

Progress in understanding uterine artery dysfunction in hypertensive pregnancies remains constrained by the inherent challenges of studying the uterine vasculature during human pregnancy. Consequently, small animal models, particularly the rat, have been instrumental for mechanistic studies due to their comparable vascular adaptations and hemochorial placentation (16,17). The spontaneously hypertensive stroke-prone rat (SHRSP), a well-established model of genetic hypertension and cardiovascular disease, provides a translatable model to study cardiovascular dysfunction in hypertensive disorders of pregnancy (18). The SHRSP was derived through selective breeding of spontaneously hypertensive rats for stroke susceptibility and shares a common Wistar origin with the normotensive Wistar Kyoto (WKY) control strain. In addition to hypertension, adult SHRSP exhibit left ventricular hypertrophy (19), insulin resistance (20), and renal damage (21). Importantly, pregnancy in the SHRSP occurs in the context of pre-existing hypertension, providing a model to investigate how chronic maternal cardiovascular disease interacts with the physiological demands of pregnancy, an increasingly prevalent and poorly understood clinical scenario. Compared to pregnant WKY rats, pregnant SHRSP exhibit reduced uterine blood flow and fetal growth restriction (22,23). While these findings establish the SHRSP as a robust model to study the effects of cardiovascular dysfunction on pregnancy, the molecular mechanisms driving uterine vascular dysfunction remain poorly defined.

To address this gap, we analyzed uterine artery structure in pregnant WKY and SHRSP at mid-pregnancy, and performed transcriptomic profiling of these vessels to identify genes and pathways associated with vascular dysfunction during hypertensive pregnancy. Through these analyses, we identified *Spp1*, encoding the matricellular protein and proinflammatory cytokine osteopontin (OPN), as markedly upregulated in uterine arteries of pregnant SHRSP. OPN is a recognized mediator of vascular remodelling and inflammation in cardiovascular disease, with elevated circulating levels in individuals with hypertension (24). In animal models, OPN promotes aortic wall thickening and macrophage infiltration (25–27), and is highly expressed in uterine natural killer (NK) cells that drive pregnancy-dependent vascular remodelling (28–30). However, the role of OPN in uterine vascular remodelling during pregnancy remains unexplored. We therefore used an OPN-deficient SHRSP model to investigate whether OPN contributes to impaired uterine vascular transformation and fetal growth restriction in hypertensive pregnancy. Our findings implicate OPN as a mediator of uterine vascular dysfunction and suggest that it contributes to the impaired vascular remodelling characteristic of hypertensive pregnancy.

## Materials and Methods

### Data Availability

Data that support the findings of this study are available from the corresponding author upon reasonable request. Large-scale datasets generated during this study have been deposited in publicly available repositories and are described in the relevant Methods sections. Additional methodological details are provided in the Supplemental Material.

### Animals

WKY rats were purchased from Charles River Laboratories (Senneville, QC, Canada). SHRSP *Spp1*^em1Mcwi^ (RRID:RGD_11553896) were obtained as heterozygotes from the PhysGen Knockout Program at the Medical College of Wisconsin (Milwaukee, WI, USA), and subsequently bred in-house to generate SHRSP wild-type (WT) and SHRSP *Spp1*(Δ/Δ) animals for experiments. Rats were maintained under a 12-hour light/dark cycle with food and water provided *ad libitum*. Females (approximately 10 weeks old) were mated with males (12-16 weeks old) of the same strain and genotype. Gestational day (GD) 0.5 was defined as the day following spermatozoa detection in a vaginal lavage. All animal procedures were approved by the Western Animal Care Committee and adhered to the guidelines of the Canadian Council on Animal Care.

### Experimental protocol and tissue collection

For fetal measurements and artery histology, pregnant WKY and SHRSP WT dams were euthanized at GD15.5 (N=6 per strain) or GD18.5 (N=4-6 per strain) using mild carbon dioxide inhalation until respiratory failure, followed by cardiac puncture and blood collection. Upon dissection, viable fetuses were counted. Uterine arteries, first order mesenteric arteries, and fetuses were collected, fixed in 10% neutral buffered formalin for 48 hours, and then transferred to 70% ethanol until further processing. Uterine arteries were not collected from uterine horns containing fewer than four fetuses. Fetal crown-rump length was measured using a digital caliper, followed by fetal weight.

For SHRSP WT versus SHRSP *Spp1*(Δ/Δ) comparisons, a separate cohort of pregnant dams were euthanized at GD15.5 (N=6 per strain) or GD18.5 (N=3-6 per strain) for uterine artery and fetal analyses.

For ultrasound and RNA sequencing, pregnant WKY, SHRSP WT, and SHRSP *Spp1*(Δ/Δ) dams underwent ultrasound imaging and were euthanized at GD15.5 (N=6 per strain). Uterine arteries were dissected, cleared of surrounding adipose tissue, snap-frozen in liquid nitrogen, and stored at -80°C until processing. To minimize animal use, SHRSP WT ultrasound and sequencing data were shared between WKY versus SHRSP WT and SHRSP WT versus SHRSP *Spp1*(Δ/Δ) comparisons.

### Artery histology and morphometry

Following fixation, tissues were dehydrated, embedded in paraffin, and sectioned at 5 𝜇m thickness. Arteries were then stained with hematoxylin and eosin and digitally scanned using a Leica Aperio AT2 Brightfield Slide Scanner. Total vessel area (defined from the lumen to the outer boundary of the tunica media; tunica adventitia excluded), lumen area, wall thickness, and vessel diameter were quantified by manual tracing and a digital ruler using Aperio ImageScope. Measurements were averaged across six serial cross-sections per artery.

### Ultrasound

Ultrasound imaging and Doppler waveform recordings were used to assess uterine artery blood flow parameters at GD15.5 using a VisualSonics Vevo 2100 preclinical ultrasound imaging system equipped with a 20 MHz MicroScan transducer (model MS-250, VisualSonics), as previously described (31). Doppler waveforms were acquired from the left and right uterine arteries near the utero-cervical junction and adjacent to a fetal-placental unit. Peak systolic velocity (PSV) and end diastolic velocity (EDV) measurements were obtained, and the resistance index ([PSV-EDV]/PSV) and systolic-to-diastolic ratio (PSV/EDV) were calculated and averaged between arteries.

### RNA extraction and quantitative RT-PCR

RNA was extracted from uterine arteries by homogenizing in TRA-zol (GeneBio Systems), followed by chloroform extraction, collection of the aqueous phase, and dilution 1:1 with 70% ethanol. Samples were then placed on RNeasy columns (Qiagen Inc.), and RNA was purified according to the manufacturer’s protocol, including DNase I treatment to remove residual genomic DNA. Complementary DNA (cDNA) was generated using a High-Capacity cDNA Reverse Transcription Kit (ThermoFisher Scientific). Samples were mixed with SensiFAST SYBR green PCR Master Mix (FroggaBio) and gene-specific primers (Table S1). Amplification and fluorescence detection were performed using a CFX96 Real-Time PCR Detection System (Bio-Rad Laboratories), with an initial denaturation at 95℃ for 10 minutes, followed by 40 cycles of 95℃ for 15 seconds and 60℃ for 1 minute. Relative gene expression was calculated using the comparative cycle threshold (ΔΔCt) method (32), using the geometric means of *Gapdh* and *Actb* as reference RNA.

### RNA sequencing

RNA was extracted from WKY, SHRSP WT, and SHRSP *Spp1*(Δ/Δ) uterine arteries as described above, and submitted to Génome Québec (Montréal, QC, Canada) for library preparation and RNA sequencing. RNA integrity was assessed using a Bioanalyzer (Agilent Technologies), with all samples achieving an RNA integrity number greater than 8.2. Libraries were generated using the NEBNEXT Ultra RNA Library Prep Kit for Illumina (New England Biolabs) and sequenced using a NovaSeq 6000 (Illumina) with 100 bp paired-end reads, generating at least 30 million reads per sample with a phred score of 39. Adapter and overrepresented sequences were removed, and reads were aligned to the mRatBN7.2 rat genome using Bowtie2. Analysis for WKY versus SHRSP WT and SHRSP WT versus SHRSP *Spp1*(Δ/Δ) were performed separately. Read counts were normalized and differential expression was performed using DESeq2 (version 1.50.2) within R (version 4.5) using the Bioconductor (version 3.22) package manager (33). For WKY versus SHRSP WT, genes with an adjusted *P*<0.01 and absolute Log_2_FoldChange>1 were considered differentially expressed. For SHRSP WT versus SHRSP *Spp1*(Δ/Δ), genes with an adjusted *P*<0.05 were considered statistically differentially expressed, with no fold-change threshold applied. Gene Ontology (GO) and gene set enrichment analysis were performed using the clusterProfiler (version 4.18.4) and enrichplot packages (version 1.30.4). The data have been deposited in the Gene Expression Omnibus database maintained by the National Center for Biotechnology Information (accession number GSE330799).

### Statistical analysis

Statistical analysis for RNA sequencing is described above. For all other analyses, data are expressed as mean ± SEM. An unpaired Student’s t-test was used for comparisons between two groups. One-way or two-way analysis of variance (ANOVA) with appropriate post-hoc tests were used for comparisons between three or more groups or for analyses involving two independent variables, respectively. A linear mixed model was used to account for litter effects where appropriate. When data were not normally distributed, a non-parametric test was used. Differences were considered statistically significant if *P*<0.05. Analyses were performed using GraphPad Prism 10.6.0 or RStudio. The number of dams (N) and fetuses (n; where relevant) used for experiments are indicated in the figure legends.

## Results

### SHRSP pregnancies exhibit hypertension and impaired fetal growth

SHRSP display chronic hypertension and metabolic dysfunction (18,20), complications that are increasingly prevalent in pregnancy. To confirm that this model recapitulates key aspects of hypertensive disorders of pregnancy, we assessed their blood pressure relative to normotensive WKY controls. As expected, SHRSP dams exhibited elevated systolic, diastolic, and mean arterial blood pressure prior to and throughout pregnancy, confirming the hypertensive phenotype (Figure 1A).

**Figure 1.**
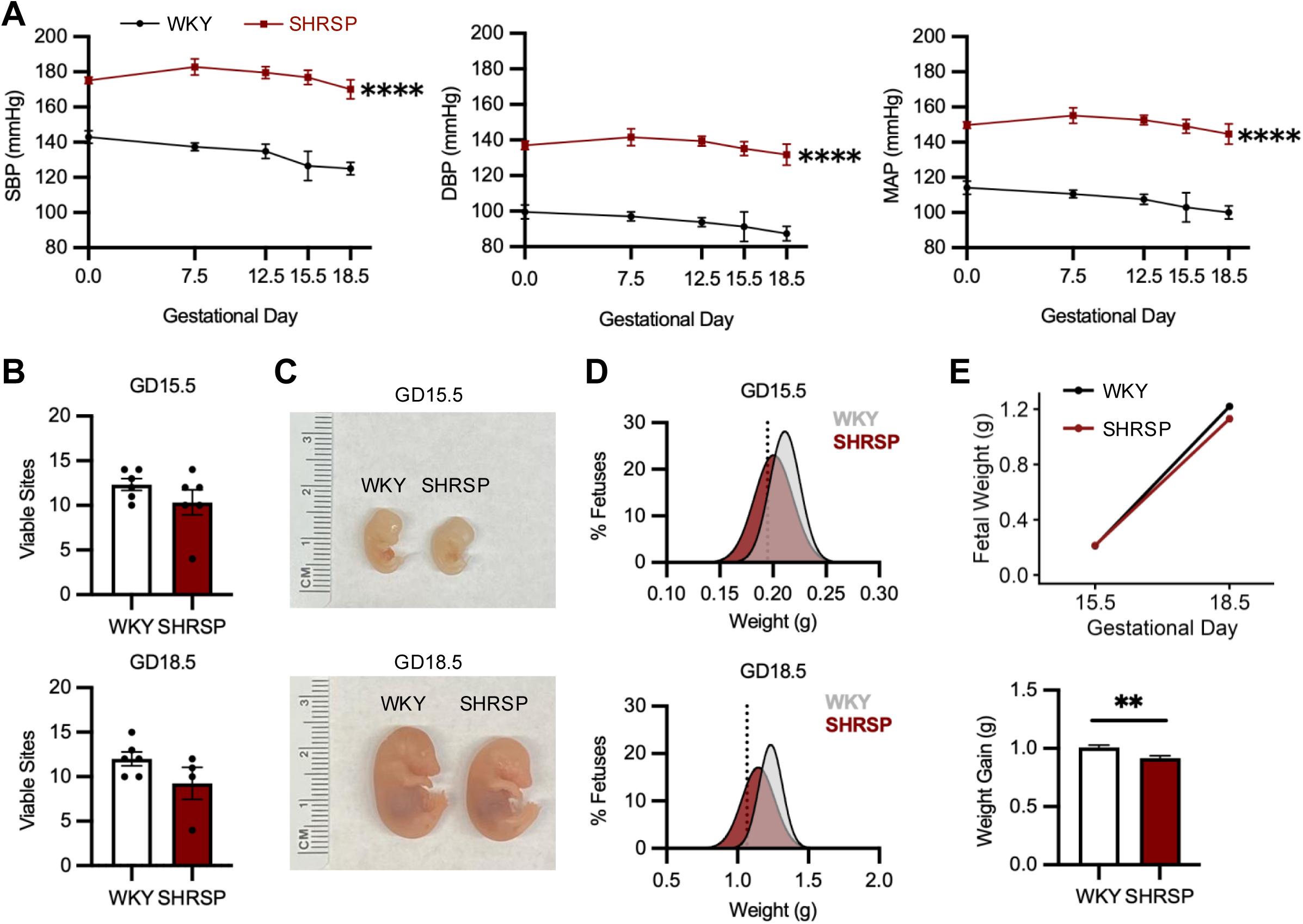
Hypertension and reduced fetal growth in SHRSP pregnancies. (**A**) Systolic, diastolic, and mean arterial pressure (SBP, DBP, MAP) across gestation (N=5). Area under the curve analysis was used for statistics. (**B**) Number of viable fetuses at GD15.5 and 18.5 (N=4-6 dams). (**C**) Representative images of GD15.5 and 18.5 fetuses. (**D**) Fetal weight distribution curves at GD15.5 (WKY n=69 fetuses, N=6 dams; SHRSP n=57 fetuses, N=6 dams) and 18.5 (WKY n=72 fetuses, N=6 dams; SHRSP n=37 fetuses, N=4 dams). Vertical dotted line indicates 10^th^ percentile of fetal weight for WKY pregnancies. (**E**) Fetal weight growth trajectory and average fetal weight gain from GD15.5 to 18.5. Mann-Whitney U test was performed for **(B)**, and a linear mixed model was used for **(E)** to account for litter effects. Asterisks denote statistical significance (^**^, *P*<0.01; ^****^, *P*<0.0001).

Fetal growth and survival were assessed at mid- and late-gestation (GD15.5 and GD18.5), timepoints that span robust uteroplacental vascular remodelling and exponential fetal growth in rats. The number of viable fetuses did not differ between strains at either timepoint (Figure 1B). However, fetuses from SHRSP pregnancies appeared visibly smaller than those from WKY (Figure 1C) and were significantly reduced in length at GD15.5 and both weight and length at GD18.5 (Figure S1). Using WKY as the reference, we next measured the proportion of fetuses that fell below the tenth percentile, which is commonly used as a proxy for fetal growth restriction (34). At GD15.5, 42% of SHRSP fetuses fell below the tenth percentile versus 9% in WKY (*P*<0.0001; Table S2, Figure 1D). A similar leftward shift in weight distribution persisted at GD18.5 (Figure 1D), although group differences were no longer statistically significant (19% SHRSP versus 10% WKY; Table S2).

To further characterize fetal growth, we assessed weight trajectories across gestation using a linear mixed-effects model with strain and GD as fixed effects, and dam as a random effect to account for litter variations. A significant strain × gestational day interaction was observed (*P*=0.00523), indicating that fetal growth differed between strains over time (Figure 1E). Specifically, SHRSP fetuses exhibited reduced weight gain from GD15.5 to GD18.5 compared to WKY (0.916 g vs. 1.008 g in WKY; Figure 1E). Together, these findings demonstrate that SHRSP pregnancies exhibit hypertension and impaired fetal growth.

### Impaired uterine artery remodelling in SHRSP pregnancies

Successful pregnancy requires extensive remodelling of the uterine arteries to support high-volume blood flow to the uterus and sustain fetal growth. In hypertensive disorders of pregnancy, this process is impaired, resulting in high-resistance vessels and reduced uterine perfusion, which are associated with fetal growth restriction (14,15). Accordingly, we characterized uterine artery structure at GD15.5 and GD18.5 in SHRSP and WKY (Figure 2A). In WKY, total vessel area increased across gestation, without changes to lumen-to-vessel area ratio or wall thickness (Figure 2B), consistent with outward hypertrophic remodelling. In contrast, in SHRSP uterine arteries, vessel area did not increase as gestation progressed, and they displayed significantly reduced lumen area and increased wall thickness at GD15.5 (Figure 2B), indicative of inward hypertrophic remodelling. SHRSP uterine arteries also exhibited many conspicuous unstained spaces within the tunica media (Figure 2A). Although the nature of these spaces remains unclear, they may reflect altered medial organization or integrity. Differences between strains were less pronounced at GD18.5 (Figure 2B), suggesting that structural abnormalities are most apparent at mid-gestation and that uterine artery remodelling is delayed in SHRSP.

**Figure 2.**
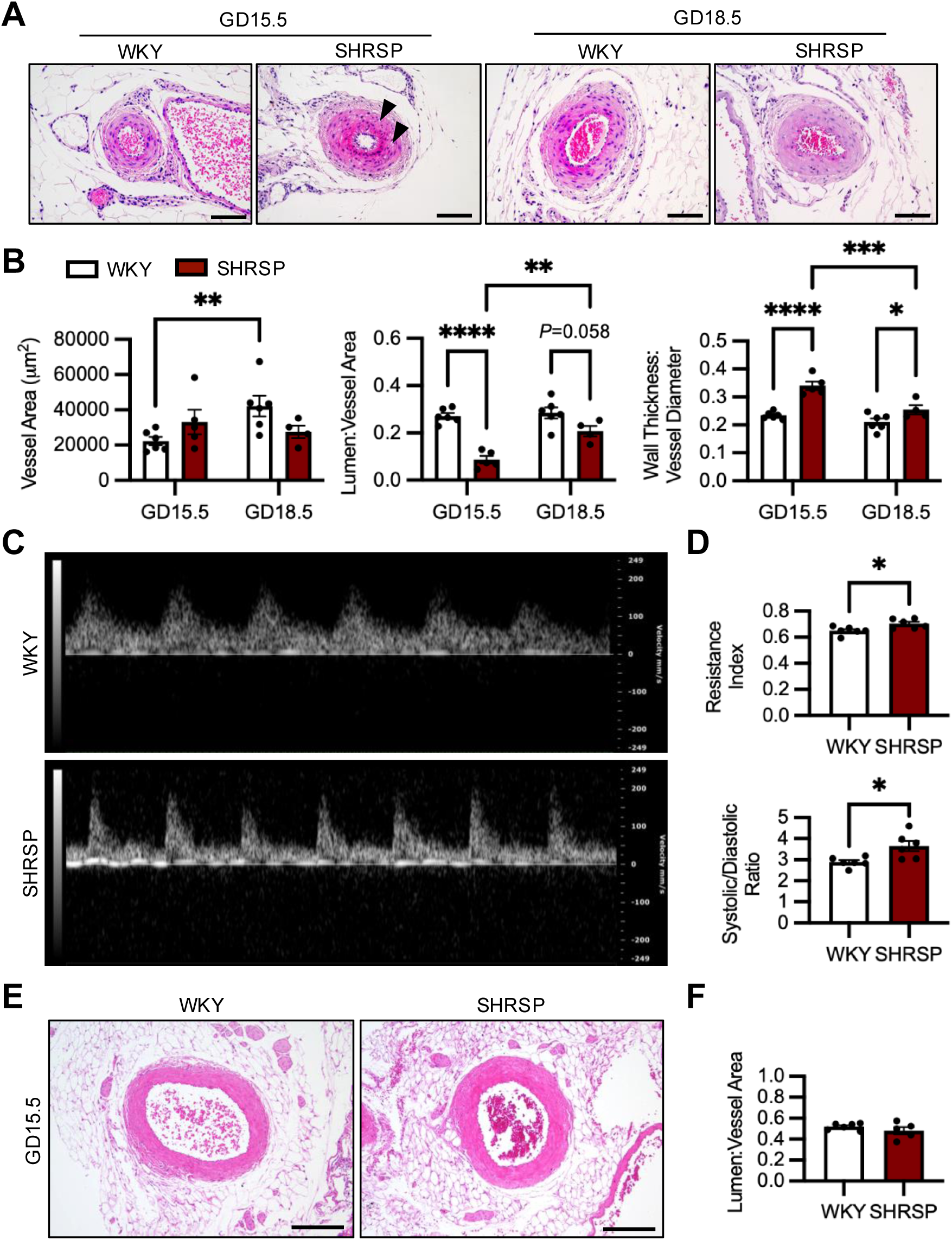
Impaired uterine artery remodelling in hypertensive pregnancy. (**A**) GD15.5 and 18.5 WKY and SHRSP uterine arteries stained with hematoxylin and eosin. Arrowheads denote unstained areas in tunica media. Scale bar; 100 µm. (**B**) Quantification of vessel area, lumen/vessel area ratio, and wall thickness/vessel diameter ratio (N=4-6). (**C**) Representative Doppler ultrasound recordings at GD15.5 over 1.3 seconds. (**D**) Resistance index and systolic/diastolic ratio of GD15.5 uterine arteries (N=6). (**E**) GD15.5 mesenteric artery stained with hematoxylin and eosin. Scale bar; 200 µm. (**F**) Quantification of lumen/vessel area ratio (N=6). Asterisks denote statistical significance (^*^, *P*<0.05; ^**^, *P*<0.01; ^***^, *P*<0.001; ^****^, *P*<0.0001) using two-way ANOVA followed by Tukey’s post hoc test for (**B**) and Student’s unpaired t-test for (**D**) and (**F**).

To determine whether these structural changes translated into functional impairment, uterine artery blood flow was assessed by Doppler ultrasound at GD15.5 (Figure S2). SHRSP uterine arteries exhibited altered Doppler waveforms (Figure 2C), resulting in increased resistance index and systolic-to-diastolic ratio compared to WKY (Figure 2D). These findings indicate elevated vascular resistance and reduced uterine perfusion in SHRSP, consistent with impaired uterine artery remodelling and its association with fetal growth restriction.

Another vessel that undergoes structural remodelling during pregnancy, albeit not to the same extent as the uterine artery, is the mesenteric artery (35). Thus, we evaluated whether the vascular dysfunction in SHRSP was specific to the uterine artery or more broadly applied to other vessels. No differences in vessel morphology or lumen-to-vessel area ratio were observed in mesenteric arteries between WKY and SHRSP on GD15.5 (Figure 2E-F), suggesting that arterial dysfunction during hypertensive pregnancy was more localized to the uterine vasculature. Thus, SHRSP pregnancies exhibit impaired uterine artery remodelling and delayed vascular adaptation to pregnancy.

### Enhanced inflammatory and tissue remodelling pathways in SHRSP uterine arteries

Impaired uterine artery remodelling in SHRSP pregnancies suggests an underlying pathological resistance to this process. To identify molecular pathways that contribute to these differences, we performed bulk RNA sequencing on uterine arteries from WKY and SHRSP at GD15.5. Principal component analysis revealed clear separation between arteries from normotensive and hypertensive dams, indicating distinct transcriptional profiles (Figure 3A). Differential expression analysis identified 302 upregulated and 364 downregulated genes in SHRSP uterine arteries relative to WKY.

**Figure 3.**
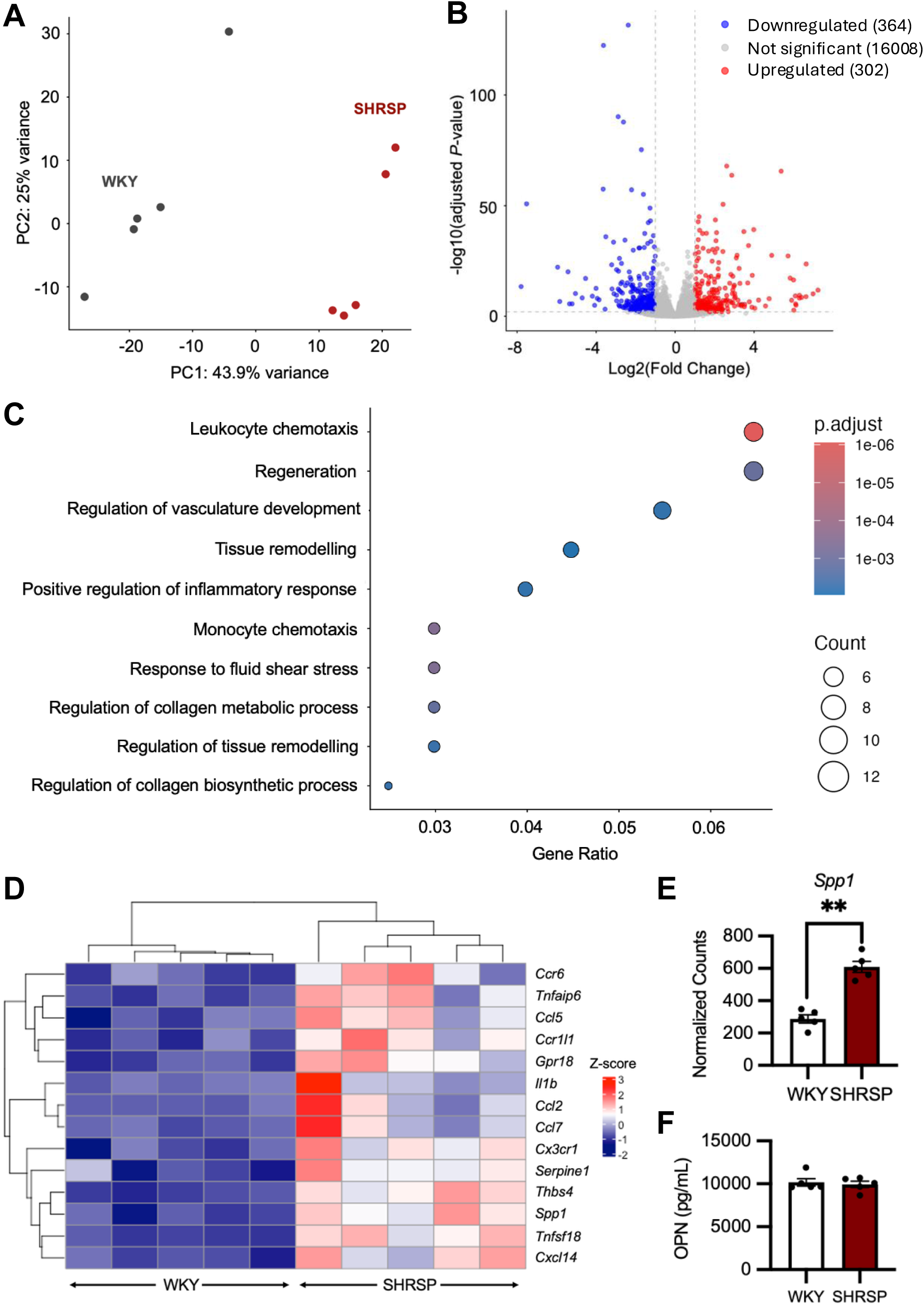
Transcriptional profiling identifies inflammatory pathway changes in uterine arteries during hypertensive pregnancy. **(A)** Principal component analysis showing sample clustering and variance (N=5). **(B)** Volcano plot depicting differentially expressed genes. Significantly upregulated genes in SHRSP are in red, downregulated genes in blue, and nonsignificant genes in grey (*P_adj_*<0.01, Log_2_FoldChange>1). **(C)** Biological process GO terms from upregulated genes in SHRSP WT compared to WKY. **(D)** Clustered heatmap for the GO term “Leukocyte chemotaxis.” **(E)** Normalized expression counts for *Spp1* (^**^, *P_adj_*<0.01). **(F)** Concentration of OPN in GD15.5 maternal blood plasma (N=5; Student’s unpaired t-test).

GO enrichment analysis revealed that the most significantly upregulated pathways in SHRSP uterine arteries were related to leukocyte chemotaxis and migration (Figure S3A), consistent with an enhanced inflammatory state. Additional enriched pathways included vascular and tissue remodelling, as well as responses to fluid shear stress (Figure 3C), aligning with the morphological and functional evidence of impaired uterine artery remodelling in SHRSP. Downregulated pathways were related to cell-cell adhesion and extracellular matrix organization (Figure S3B), indicating compromised vessel integrity.

To further examine genes associated with enhanced inflammation in SHRSP uterine arteries, we visualized genes within the “leukocyte chemotaxis” GO term (Figure 3D). Several genes implicated in inflammation and vascular dysfunction, including *Ccl5*, *Il1b*, *Ccl2*, and *Serpine1*, were upregulated in SHRSP. Notably, *Spp1*, encoding the matricellular protein and proinflammatory cytokine OPN, was among the most highly upregulated genes in this pathway. It also featured across multiple upregulated pathways, including responses to fluid shear stress and tissue remodelling (Figure 3D-E; Table S3), underscoring its potential importance in uterine artery dysfunction. To determine whether increased *Spp1* expression in the uterine vasculature translated to elevated circulating OPN, maternal plasma levels were measured at GD15.5 by ELISA. No differences were observed between strains (Figure 3F), consistent with the notion that changes are localized to the uterine vasculature. Collectively, these data implicate OPN as a potential mediator of inflammatory and structural changes in the uterine artery during hypertensive pregnancy.

### OPN deficiency improves uterine artery remodelling and fetal growth trajectory in SHRSP

Given the potential role of OPN in driving uterine artery dysfunction during hypertensive pregnancy, we characterized a novel SHRSP strain harboring a 5 bp deletion in coding exon 4 of *Spp1* (Figure 4A). SHRSP containing the WT *Spp1* allele and/or the *Spp1***Δ** allele was determined using a PCR-based genotyping strategy (Figure 4B-C). The mutation introduced a frameshift and caused a predicted premature termination codon (Figure 4D). Because *Spp1* is highly expressed in the kidney (36), we assessed its expression in this tissue to confirm loss in *Spp1-*mutant animals (Figure 4E). Consistent with this, OPN was undetectable in the kidneys of *Spp1*-mutant rats by immunofluorescence and in maternal plasma on GD15.5 (Figure 4E-F). Together, these findings confirm that *Spp1-*mutant animals are deficient in OPN.

**Figure 4.**
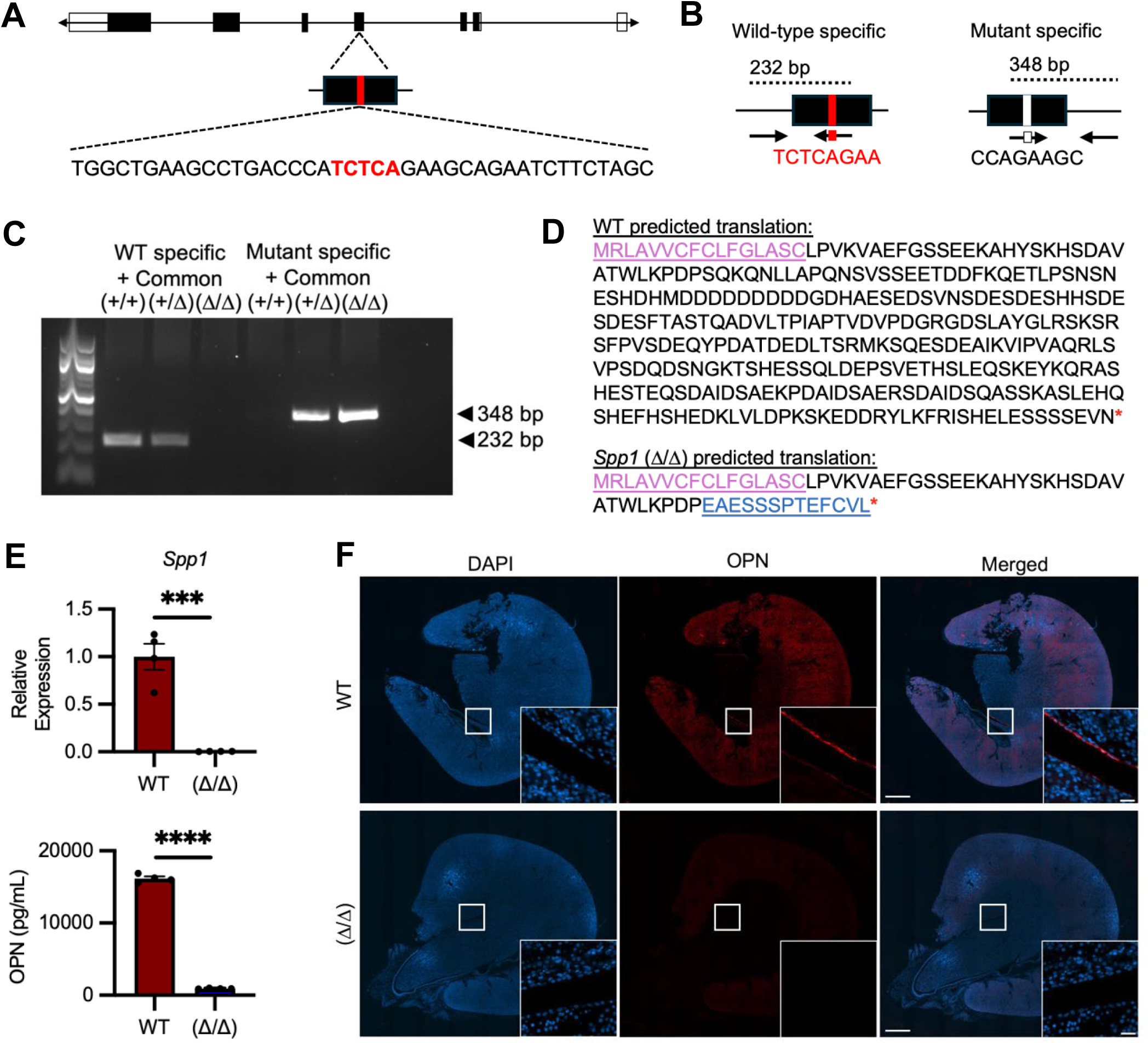
SHRSP *Spp1*(Δ/Δ) are OPN-deficient. **(A)** Mutation target site in the fourth coding exon of *Spp1.* **(B)** Genotyping strategy used to identify rats containing the WT *Spp1* allele (+) and/or the *Spp1***Δ** allele. **(C)** Agarose gel electrophoresis of PCR products. 232 bp band represents WT *Spp1* allele and 348 bp bands represents *Spp1***Δ** allele. **(D)** Predicted amino acid sequence encoded by WT and *Spp1***Δ** (pink, leader peptide; blue, deviations from WT sequence; red asterisk, stop codon). **(E)** Relative *Spp1* expression in GD15.5 maternal kidney and concentration of OPN in GD15.5 maternal blood plasma (N=4). **(F)** OPN localization to GD15.5 kidney cortex and calyx region (inset) in SHRSP WT rats and absence in *Spp1***Δ/Δ** rats. Scale bar; 1000 µm and 50 µm (inset). Asterisks denote statistical significance (^***^, *P*<0.001; ^****^, *P*<0.0001) using Student’s unpaired t-test.

Given prior evidence linking OPN to vascular dysfunction in cardiovascular disease, we first assessed whether OPN deficiency alters the hypertensive phenotype in SHRSP. No differences in systolic, diastolic, or mean arterial blood pressure were observed before and during gestation (Figure S4), indicating that OPN deficiency does not affect systemic blood pressure.

Subsequent analyses focused on the uterine artery, given our findings of localized vascular remodelling defects in this vessel in SHRSP. At GD15.5, OPN-deficient SHRSP exhibited improved uterine artery structure, as shown by an increased lumen-to-vessel area ratio compared to SHRSP WT, without a change in wall thickness (Figure 5A-B). To determine whether these structural changes translated into functional improvement, uterine artery blood flow was assessed by Doppler ultrasound at GD15.5. OPN-deficient SHRSP displayed reduced vascular resistance (Figure 5C), evidenced by decreased resistance index and systolic-to-diastolic ratio relative to SHRSP WT (Figure 5D). Therefore, loss of OPN improves uterine artery structure and function without altering systemic blood pressure, supporting a role for OPN in mediating uterine vascular dysfunction during hypertensive pregnancy.

**Figure 5.**
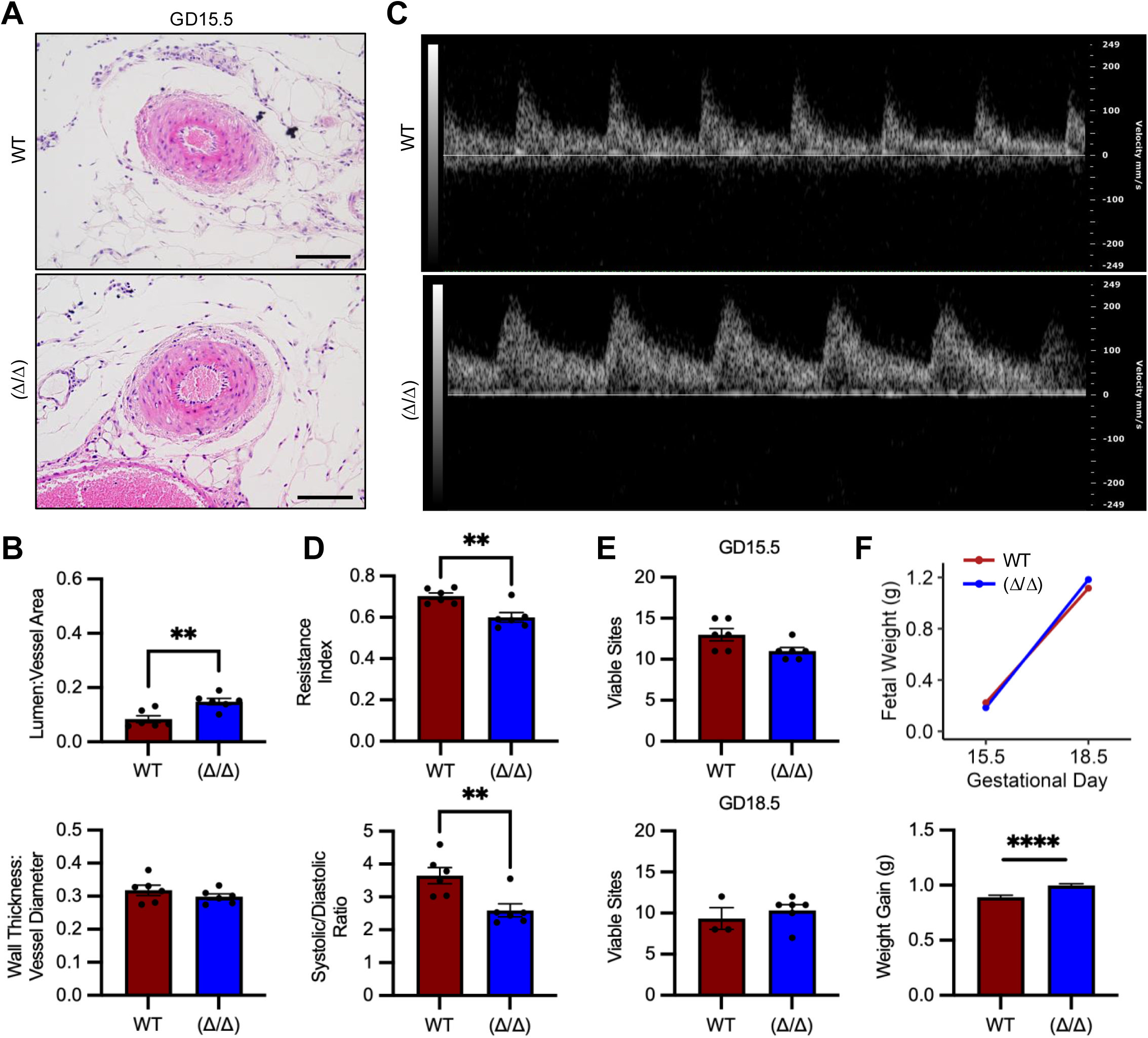
OPN deficiency improves uterine artery structure and function and enhances fetal growth trajectory in SHRSP pregnancy. **(A)** GD15.5 SHRSP WT and *Spp1*(Δ/Δ) uterine arteries stained with hematoxylin and eosin. Scale bar; 100 µm. **(B)** Quantification of lumen/vessel area ratio and wall thickness/vessel diameter ratio (N=6). **(C)** Representative Doppler ultrasound recordings at GD15.5 over 1.3 seconds. **(D)** Resistance index and systolic/diastolic ratio of GD15.5 uterine arteries (N=6). **(E)** Number of viable fetuses from GD15.5 and 18.5 pregnancies (N=3-6 dams each). **(F)** Fetal weight trajectory and average fetal weight gain from GD15.5 to 18.5. Student’s unpaired t-test was performed for (**B**) and (**D**), Mann-Whitney U test for (**E**), and a linear mixed model for (**F**) to account for potential litter effects. Asterisks denote statistical significance (^**^, *P*<0.01; ^****^, *P*<0.0001).

Decreased uterine artery resistance index in the OPN-deficient SHRSP pregnancies suggests increased perfusion toward the placenta. To determine whether this was associated with improved fetal outcomes, we assessed fetal viability and growth at GD15.5 and GD18.5. Although OPN deficiency did not affect the number of viable fetuses at either gestational timepoint (Figure 5E), analysis of fetal weight trajectory using a linear mixed-effects model revealed a significant strain × gestational day interaction (*P*=9.94×10^-7^), indicating that fetal growth trajectories differed between groups over time. Specifically, OPN-deficient SHRSP fetuses exhibited a greater weight gain from GD15.5 to GD18.5 compared to SHRSP WT (0.999 g vs. 0.892 g; Figure 5F). At GD18.5, a greater proportion of OPN-deficient fetuses trended toward the upper end of the distribution (18% OPN-deficient fetuses vs. 7% SHRSP WT fetuses above the 90^th^ percentile; Table S4). Together, these findings suggest that improved uterine artery function in OPN-deficient SHRSP pregnancies may enhance fetal growth by supporting greater uteroplacental perfusion.

### OPN deficiency does not alter uterine NK cell recruitment

In pregnancy, NK cells are abundant within the uterus and play key roles in the pregnancy-dependent transformation of spiral arteries – downstream branches of the uterine arteries – which promote adequate blood flow to the placenta to sustain fetal growth. OPN has been implicated in the maintenance of peripheral NK cell populations (37), and is strongly expressed by uterine NK cells (28–30). Therefore, we investigated whether loss of OPN alters uterine NK cell abundance as a potential mechanism underlying uterine vascular maladaptation.

We first assessed uterine NK cell number during early gestation (GD9.5), when NK cells are abundant in the uterus, by staining for the pore-forming protein perforin. Consistent with prior reports, OPN staining overlapped with perforin, indicating that OPN is expressed by uterine NK cells. However, no differences in uterine NK cell number were observed between OPN-deficient and SHRSP WT (Figure S5A-B). Similarly, expression of genes associated with NK cell maturation and function were unchanged by OPN deficiency (Figure S5C). At mid-gestation, uterine NK cells remained abundant around spiral arteries in both WT and OPN-deficient rats, indicating that loss of OPN does not affect NK cell abundance or spatial distribution. Strikingly, less uterine NK cells appeared to co-localize with OPN at GD15.5 compared to GD9.5 (Figure S6). These findings suggest that improved uterine vascular function in OPN-deficient rats is likely independent of effects on NK cell recruitment.

### Loss of OPN partially restores a healthy vascular gene signature in SHRSP uterine arteries

Given the improved uterine artery function in OPN-deficient SHRSP, we next assessed whether the inflammatory transcriptional profile characteristic of SHRSP uterine arteries was diminished. Differential expression analysis identified 20 downregulated and 11 upregulated genes in OPN-deficient uterine arteries relative to SHRSP WT controls (Figure 6A). Notably, OPN deficiency partially normalized the SHRSP uterine artery transcriptional profile, with several genes upregulated in SHRSP WT restored toward WKY levels (Figure 6B). Gene set enrichment analysis further demonstrated reduced enrichment of “cell chemotaxis” and “cell surface receptor signalling pathway via JAK-STAT” in OPN-deficient SHRSP relative to WT (Figure 6C). These pathways are implicated in vascular smooth muscle cell proliferation and immune cell recruitment, processes that drive pathological vessel remodelling (38).

**Figure 6.**
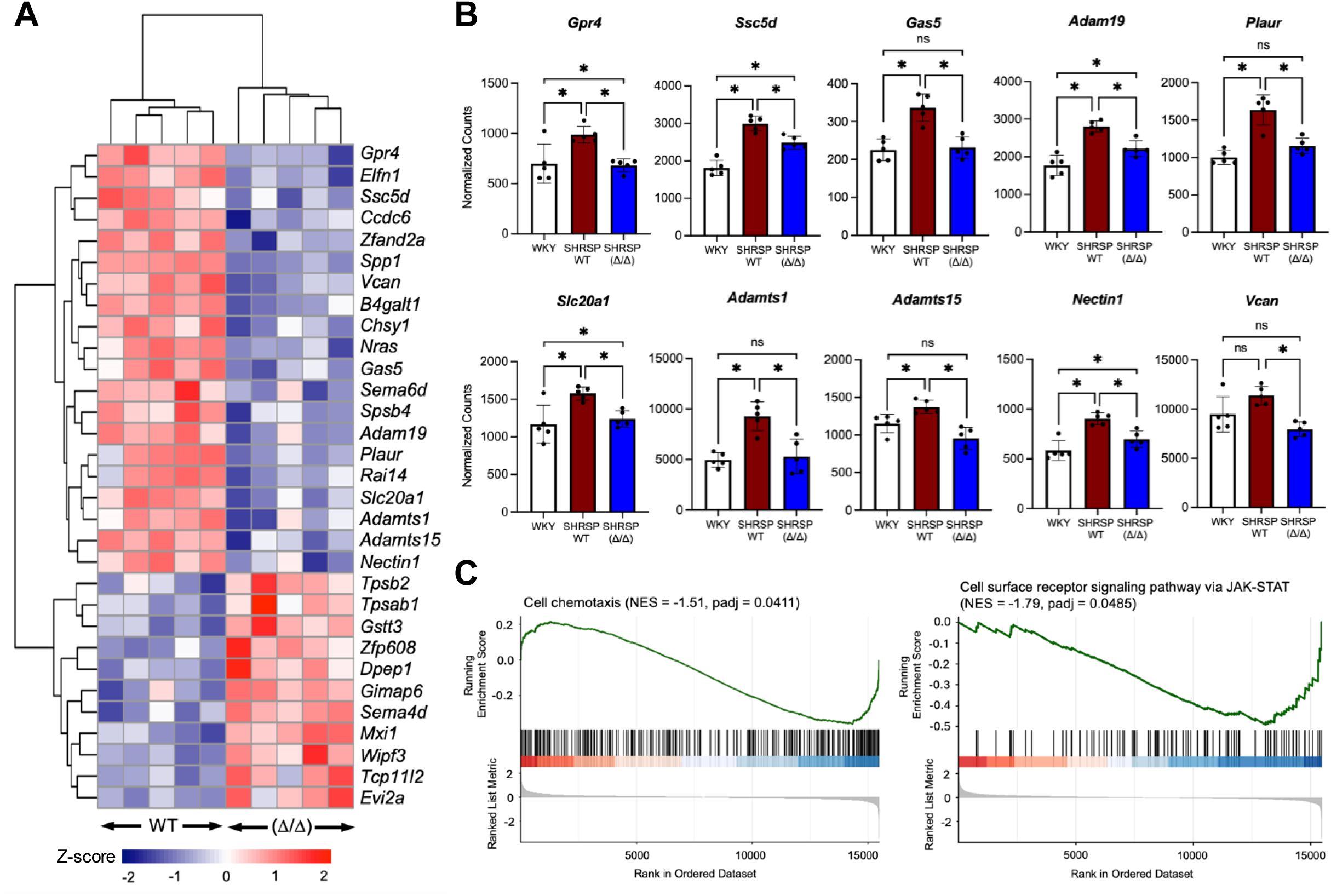
OPN deficiency attenuates inflammatory and vascular dysfunction-associated gene expression in uterine arteries from hypertensive pregnancies. (A) Clustered heat map of differentially expressed genes in SHRSP *Spp1*(Δ/Δ) compared to WT (*P_adj_*<0.05). (B) Select genes upregulated in hypertensive pregnancy (SHRSP WT versus WKY) and reversed by OPN deficiency (^*^, *P_adj_*<0.05). (C) Gene set enrichment analysis for the ontology terms “Cell chemotaxis” and “Cell surface receptor signalling pathway via JAK-STAT.” Differentially expressed genes are denoted by vertical black lines

## Discussion

This study used the chronically hypertensive SHRSP model to uncover mechanisms underlying uterine artery dysfunction during hypertensive pregnancy. We found that pregnant SHRSP exhibit impaired uterine artery adaptation and reduced fetal growth, accompanied by increased expression of inflammatory and vascular pathology-associated genes. Among these, *Spp1*, which encodes OPN, emerged as a candidate mediator of vascular dysfunction. Using a novel OPN-deficient SHRSP model, we demonstrate that loss of OPN restores uterine artery morphology and blood flow, attenuates inflammation, and improves fetal growth trajectory. Together, these findings identify OPN as a contributor to uterine artery dysfunction in hypertensive pregnancy.

The SHRSP provides a translationally relevant model of chronic hypertension that captures clinically important features of human hypertensive disease and enables mechanistic investigation of impaired uteroplacental vascular adaptation during pregnancy. A key strength of this model is that hypertension develops spontaneously, without genetic, surgical, or pharmacological manipulation, paralleling the idiopathic nature of most human hypertension (39). Consistent with fetal growth patterns observed in human hypertensive pregnancies (40), SHRSP fetuses were modestly smaller than WKY fetuses, with a greater proportion falling below the 10^th^ percentile. Although reduced fetal growth in SHRSP pregnancies has not been uniformly reported across studies (22,23), these discrepancies may reflect differences in analytical approaches, including whether fetal growth was assessed at a single gestational age or across multiple time points, and how litter effects were handled. Averaging fetal weight within a litter may obscure biologically meaningful variation and underestimate the extent of impaired fetal growth.

Our findings extend previous work showing uterine artery dysfunction in SHRSP pregnancy (22). In normotensive WKY pregnancies, uterine arteries undergo outward hypertrophic growth, consistent with vascular adaptation in healthy pregnancy (12). In contrast, SHRSP uterine arteries displayed inward hypertrophic remodelling, including reduced lumen area, increased wall thickness, elevated resistance index, and reduced blood flow at GD15.5. This demonstrates that impaired uterine artery adaptation is already evident by mid-gestation, even earlier than what was previously reported (22).

To our knowledge, this study is the first to characterize the uterine artery transcriptome at mid-gestation in a model of hypertensive pregnancy. We identified an enrichment of leukocyte chemotaxis and inflammatory signalling in SHRSP uterine arteries. These findings are consistent with previous work showing increased immune activation in SHRSP uterine arteries at GD6.5, before overt structural or functional differences are apparent (41). Notably, several genes, including *Ccl7, Cxcl14, Ccr1* and *Il6,* were elevated at both GD6.5 and GD15.5 across studies. Together, these data suggest that the SHRSP uterine artery may be primed for dysfunction early in pregnancy, with inflammatory signals preceding and potentially contributing to structural and functional abnormalities later in pregnancy.

OPN is a multifunctional matricellular protein and proinflammatory cytokine with established roles in vascular inflammation, remodelling, and fluid shear stress (42). Therefore, *Spp1* upregulation may reflect increased inflammation and altered hemodynamics in the uterine vasculature during hypertension. To test the role of OPN in uterine artery dysfunction, we used a novel OPN-deficient SHRSP model. To our knowledge, this is the first rat model combining OPN deficiency with a background of chronic hypertension. OPN-deficient SHRSP were viable and capable of sustaining pregnancy, enabling *in vivo* assessment of OPN function in hypertensive pregnancy. OPN deficiency did not alter systemic blood pressure before or during pregnancy. Despite this, OPN-deficient SHRSP exhibited larger uterine lumen areas, reduced resistance index, and improved fetal growth trajectory compared with SHRSP controls. Our findings of localized vascular improvement are consistent with a previous study in angiotensin II-infused mice, where loss of OPN protected against medial hypertrophy and inflammation in the aorta despite no change in systolic blood pressure (25). Together, these findings suggest that OPN can drive local vascular dysfunction and inflammation in hypertensive disease independent of overt changes in systemic blood pressure.

The improved vascular phenotype in OPN-deficient SHRSP was accompanied by reduced expression of inflammation-associated genes and decreased enrichment of cell chemotaxis and JAK-STAT signalling pathways. Several genes elevated in SHRSP uterine arteries were restored toward WKY levels following OPN deficiency, including *Plaur*, *Adamts1*, *Adamts15*, *Vcan*, and *Gas5*. *Plaur*, which encodes the urokinase-type plasminogen activator receptor, has been implicated in leukocyte recruitment, inflammation, and JAK-STAT signalling (43,44). The ADAMTS proteases are involved in matrix turnover, suggesting that OPN may also contribute to maladaptive matrix remodelling. Increased versican (encoded by *Vcan*) is a recognized feature of cardiovascular disease and vascular pathology (45). The concurrent upregulation of *Adamts1*, *Adamts15*, and *Vcan* in SHRSP WT arteries may reflect increased matrix remodelling activity associated with pathological vascular structure. Although the role of *Gas5* is less clear, its restoration toward WKY levels further supports a broader normalization of vascular stress-associated transcriptional programs in OPN-deficient SHRSP (46,47).

While our findings strongly support a role for OPN in uterine artery dysfunction, several limitations should be noted. First, although improved uterine artery structure and reduced resistance provide a plausible explanation for the improved fetal growth trajectory, changes in downstream vascular beds may also contribute. Spiral arteries undergo extensive remodelling during pregnancy and are essential for placental perfusion. OPN has been implicated in uterine NK cell function, trophoblast invasion, and trophoblast-endothelial cell interactions, all of which may influence spiral artery remodelling (28,48). While we found that OPN deficiency did not alter uterine NK cell recruitment to the spiral arteries, future studies could further investigate whether OPN influences spiral artery remodelling or placental vascular development in hypertensive pregnancy.

A second limitation of this study is the use of WKY rats as normotensive controls. Although WKY rats are traditionally used as controls for SHRSP in cardiovascular and pregnancy research, the strains have diverged genetically over time (49). Therefore, some differences between SHRSP and WKY uterine arteries may reflect inherent strain differences rather than hypertension-associated pathology. However, major structural differences in the mesenteric arteries were not apparent between strains, reducing the likelihood that systemic vascular differences explain the uterine artery phenotype. In addition, the improved uterine artery structure observed in OPN-deficient SHRSP supports a role for OPN in uterine artery dysfunction. Circulating maternal plasma OPN concentrations did not differ between WKY and SHRSP pregnancies, suggesting that increased *Spp1* expression mediates localized changes within the uterine vasculature rather than systemic strain-dependent differences in OPN abundance.

### Perspectives

Collectively, our findings identify OPN as a contributor to impaired uterine artery adaptation in hypertensive pregnancy. We propose that chronic hypertension creates a proinflammatory uterine vascular environment that disrupts the arterial remodelling required for expansion during pregnancy. OPN deficiency improved uterine artery structure and function in SHRSP, reduced inflammatory and vascular pathology-associated transcriptional signatures, and was associated with improved fetal growth trajectory. These findings position OPN-associated vascular inflammation as a potential mechanism linking chronic hypertension to impaired uteroplacental vascular adaptation and fetal growth restriction.

### Novelty and Relevance

#### What Is New?

This study is the first to characterize mid-gestation uterine artery dysfunction in a translatable rat model of hypertensive pregnancy, identifying inflammatory and matrix remodelling pathways associated with vascular pathology. We further identify OPN as a candidate mediator of uterine artery dysfunction and characterize a novel OPN-deficient SHRSP model that demonstrates improved uterine artery adaptation and fetal growth trajectory during hypertensive pregnancy.

#### What is Relevant?

Impaired uterine artery remodelling is a hallmark of hypertensive pregnancy and contributes to reduced uteroplacental perfusion and fetal growth restriction. Our findings identify OPN as a potential link between chronic hypertension, inflammatory signalling, extracellular matrix remodelling, and maladaptive uterine vascular adaptation.

#### Clinical/Pathophysiological Implications?

These findings support a role for OPN in the pathogenesis of uterine vascular dysfunction and suggest that OPN-associated inflammatory and remodelling pathways may represent therapeutic targets to improve vascular adaptation and fetal outcomes in hypertensive pregnancy.

## Acknowledgements

The authors gratefully acknowledge the PhysGen Knockout Program for generating the OPN-deficient rat strain used in this study and the Rat Genome Database for facilitating access to the model. We would also like to thank the Centre d’expertise et de services Génome Québec for library preparation and Karen Nygard at Biotron Integrated Microscopy Facility (Western University).

## Author Contributions

S.J.R, M.L.L designed the study and conceived of all experiments. M.L.L, J.J., V.S.P., and S.L performed experiments. Animal experiments and study design were supported by M.M. and D.G. J.C.L. and G.E.E. provided technical support and assisted with data analysis. The manuscript was written by M.L.L. and S.J.R. The manuscript was approved by all co-authors.

## Sources of Funding

This work was supported by the Canadian Institutes of Health Research (CIHR, PJT180483) to S.J.R. M.L.L was supported by a Children’s Health Research Institute Fellowship and Mitacs Accelerate grant. V.S.P was supported by NSERC Doctoral Canada Graduate Scholarship and Children’s Health Research Institute Fellowship. S.Z.L was supported by a CIHR Masters Canada Graduate Scholarship.

## Disclosures

None.

## Supplemental Material

Supplemental Methods

Tables S1-S4

Figures S1-S6

## Non-standard Abbreviations and Acronyms

DBP: Diastolic Blood Pressure
EDV: End Diastolic Velocity
GD: Gestational Day
MAP: Mean Arterial Pressure
NK: Natural Killer
OPN: Osteopontin
PSV: Peak Systolic Velocity
SBP: Systolic Blood Pressure
SHRSP: Spontaneously-Hypertensive Stroke-Prone Rat
*Spp1*: Secreted Phosphoprotein 1
WKY: Wistar Kyoto

## Notes

### Competing Interest Statement

The authors have declared no competing interest.

### Summary of Updates

Text added below will post on bioRxiv with the revised paper; Supplemental files updated.

