## Supplemental Information for "Osteopontin Mediates Uterine Artery Dysfunction in Hypertensive Pregnancy"

### Supplemental Material

#### Supplemental Methods

*Blood pressure*

Prior to breeding, virgin female WKY, SHRSP WT, and SHRSP *Spp1*(Δ/Δ) (approximately 8-weeks-old, N=5 per strain) underwent systemic blood pressure measurements using a CODA high-throughput non-invasive blood pressure system (Kent Scientific; (50)). To minimize distress during measurements, rats were acclimatized to the apparatus on three separate occasions over two weeks, and all measurements were performed by the same operator. In the third week, blood pressure measurements were recorded for virgin rats, after which the rats were mated, and measurements were repeated at GD7.5, 12.5, 15.5, and 18.5. All measurements were obtained in the morning, with 5 preliminary cuff inflations for acclimatization followed by 20 measurement cycles. Systolic blood pressure (SBP), diastolic blood pressure (DBP), and mean arterial pressure (MAP = 1/3[SBP–DBP] + DBP) were averaged across cycles. Outliers (±2 SD from the mean) were excluded, and averages were recalculated using the remaining measurements (minimum of 10 values). To minimize animal use, SHRSP WT blood pressure data were shared between WKY versus SHRSP WT and SHRSP WT versus SHRSP *Spp1*(Δ/Δ) comparisons.

*Genotyping*

A PCR-based strategy was used to determine the *Spp1* genotype. Genomic DNA was extracted from pup ear punches using reagents from the REDExtract-N-Amp DNA isolation kit (Sigma-Aldrich). DNA was amplified by PCR using Q5 High Fidelity DNA Polymerase (New England Biolabs) with WT-specific, mutant-specific, and common primers for *Spp1* (Table S1).

Thermocycler conditions included an initial denaturation step (98ºC for 1 minute), followed by 30 cycles of three-step PCR (98ºC for 10 seconds, 66ºC for 30 seconds, and 72ºC for 30 seconds), and a final extension step (72ºC for 5 minutes). PCR products were resolved on an agarose gel containing GelRed nucleic acid stain. The presence of a 232 bp band corresponded to the WT *Spp1* allele, while a 348 bp band indicated the *Spp1*Δ allele.

*Enzyme-linked immunosorbent assay (ELISA)*

Uncoagulated maternal blood was collected during dissections in an ethylenediaminetetraacetic acid-coated BD vacutainer tube, centrifuged for 10 minutes at 4°C and 1,000 × g to isolate plasma, and then frozen at -80°C until analysis. OPN levels in maternal plasma were measured using a Mouse/Rat OPN Quantikine ELISA kit (MOST00, R&D Systems) using a protocol provided by the manufacturer. A standard curve was generated by plotting absorbance values against known concentrations of recombinant OPN, and sample concentrations were determined using this curve. The sensitivity of the assay for rat OPN detection was 8.5 pg/mL.

*Immunofluorescence*

Tissues were fixed in 10% neutral buffered formalin, paraffin-embedded, and sectioned at 5 $\mu$m thickness. Sections were deparaffinized in Histo-Clear and rehydrated using decreasing concentrations of ethanol. Antigen retrieval was performed to reverse formaldehyde crosslinks by immersing sections in citrate buffer at 95°C for 20 minutes. Sections were permeabilized using 0.3% Triton X-100 and 1% bovine serum albumin in PBS, and blocked in 10% normal goat serum. Sections were then incubated overnight at 4°C with a primary antibody specific for OPN (sc-21742; 1:400; Santa Cruz Biotechnology) or perforin (TP251; 1:400; Torrey Pines Biolabs).

Mouse $IgG_1$ (5415; 1:400; Cell Signaling Technology) and rabbit IgG (2729S; 1:400; Cell Signaling Technology) were used as negative controls for OPN and perforin, respectively. The following day after washing, sections were incubated with either Alexa Fluor 555-conjugated goat anti-mouse secondary antibody (A21424, 1:250, Invitrogen), Alexa Fluor 488-conjugated goat anti-mouse secondary antibody (A11001; 1:250, Invitrogen), or Alexa Fluor 488-conjugated goat anti-rabbit secondary antibody (A11034; 1:250, Invitrogen). Nuclei were counterstained with 4′,6-diamidino-2-phenylindole (DAPI; ThermoFisher Scientific). Images were acquired using a Nikon inverted Ti2e spinning disk confocal microscope (Nikon Corporation).

*Immunohistochemistry*

Sections were deparaffinized, rehydrated, and subjected to antigen retrieval as described above. Endogenous peroxidases were blocked by immersing slides in a solution containing 0.3% $H_2O_2$ in methanol, permeabilized using 0.3% Triton X-100 and 1% bovine serum albumin in PBS, and blocked in 10% normal goat serum. Sections were then immersed in primary antibody specific for perforin (TP251; 1:400; Torrey Pines Biolabs), or rabbit IgG (2729S; 1:400; Cell Signaling Technology) as a negative control at 4°C overnight. The following day after washing, sections were immersed sequentially in biotin-conjugated secondary antibody (Sigma-Aldrich), Vectastain ABC Elite reagent (Vector Laboratories), and 3-amino-9-ethylcarbazole red (ThermoFisher Scientific). Sections were counterstained with Gill No. 1 hematoxylin (Sigma-Aldrich) and mounted using Fluoromount-G medium (ThermoFisher Scientific). Images were acquired using a Nikon inverted Ti2e spinning disk confocal microscope (Nikon Corporation).

#### Supplemental Tables and Figures

**Table S1. Forward and reverse primer sequences used for quantitative RT-PCR and genotyping.**

| Gene | Accession No. | Forward: Sequence (5’-3’) | Reverse: Sequence (5’-3’) |
|---|---|---|---|
| *Gapdh* | NM_017008.4 | GACATGCCGCCTGGAGAAAC | AGCCCAGGATGCCCTTTAGT |
| *Actb* | NM_031144 | AGCCATGTACGTAGCCATCC | CTCTCAGCTGTGGTGGTGAA |
| *Spp1* | NM_012881 | GGAGGAGAAGGCGCATTACA | TCGTCGTCGTCATCATCGTC |
| *Prf* | NM_017330 | CCCAGTGAACACAGGGAAGT | CTTCCGGTTTAGGGTTCACA |
| *Eomes* | NM_001427393 | TTCACCCAGAATCTCCCAAC | TGGAAGGCTCATTCAAGTCC |
| *Ccl5* | NM_031116 | CCTTGCAGTCGTCTTTGTCA | GAGTAGGGGGTTGCTCAGTG |
| *Vegfa* | NM_001110333 | GCAATGATGAAGCCCTGGAG | GCTCTGAACAAGGCTCACAG |
| *Tnf* | NM_012675 | CGTCGTAGCAAACCACCAAG | GAGGCTGACTTTCTCCTGGT |
| *Ogn* | NM_001106103 | CCTGCTATTGTTCGTGCCTC | TTGGTGGGCACTGATGGTAT |
| *Ptn* | NM_017066 | TCTGACTGTGGAGAATGGCA | AAGGCGGTATTGAGGTCACA |
| **For *Spp1* genotyping:** | | | |
| Wild-type specific | | | GGCTAGAAGATTCTGCTTCTGAGA |
| Mutant-specific | | CTGAAGCCTGACCCAGAAGC | |
| Common forward | | AGAAACTCAGAGACATAGAGGCA | |
| Common reverse | | | AAGCAAGAAGACCCACCACA |

**Table S2. Distribution of fetal weights from WKY and SHRSP litters at GD15.5 and 18.5.** Fetuses were stratified into percentile weight categories using the WKY fetal distribution as reference. Values are presented as number of fetuses and percentage of total fetuses per group. The proportion of growth restricted fetuses (< $10^{th}$ percentile) was compared between groups using Fisher's exact test.

| | GD 15.5 | | | GD 18.5 | | |
|---|---|---|---|---|---|---|
| | WKY | SHRSP | *P* value | WKY | SHRSP | *P* value |
| Number of Dams | 6 | 6 | | 6 | 4 | |
| Total Fetuses | 69 | 57 | | 72 | 37 | |
| < $10^{th}$ Percentile | 8.7% (6) | 42% (24) | <0.0001 | 9.7% (7) | 18.9% (7) | 0.2277 |
| 10-$90^{th}$ Percentile | 82.6% (57) | 53% (33) | | 80.6% (58) | 78.4% (29) | |
| > $90^{th}$ Percentile | 8.7% (6) | 0% (0) | | 9.7% (7) | 2.7% (1) | |

**Table S3. Biological process GO terms shown in Figure 3C and their p-adjusted value and gene ID list.** *Spp1* is marked in bold.

| GO ID | Description | *P*-adjusted | Gene ID |
|---|---|---|---|
| GO:0030595 | Leukocyte chemotaxis | 9.14E-06 | *Ccr6/Thbs4/Tnfaip6/Il1b/Cx3cr1/Ccr1l1/Ccl2/Ccl7/Ccl5/Serpine1/Tnfsf18/**Spp1**/Gpr18/Cxcl14* |
| GO:0002548 | Monocyte chemotaxis | 3.65E-04 | *Cx3cr1/Ccr1l1/Ccl2/Ccl5/Serpine1/Tnfsf18* |
| GO:0034405 | Response to fluid shear stress | 4.15E-04 | *Abca1/Gpr68/Has2/Ace/Serpine1/**Spp1*** |
| GO:0031099 | Regeneration | 1.21E-03 | *Il6/Chl1/Tnc/Ncam1/Gnat1/Runx2/Ccl2/Ace/Serpine1/**Spp1**/Afp/Lif/Prps2* |
| GO:0010712 | Regulation of collagen metabolic process | 1.43E-03 | *Fap/Il6/Ciita/Ccl2/Serpine1/Inhba* |
| GO:0034103 | Regulation of tissue remodeling | 5.76E-03 | *Ceacam1/Thbs4/Il6/**Spp1**/Siglec15/Cd24* |
| GO:0032965 | Regulation of collagen biosynthetic process | 6.26E-03 | *Il6/Ciita/Ccl2/Serpine1/Inhba* |
| GO:1901342 | Regulation of vasculature development | 6.84E-03 | *Dll1/Ceacam1/Thbs4/Il1b/Crhr2/Cx3cr1/Ccl2/Ccl5/Serpine1/Chi3l1/Lif* |
| GO:0050729 | Positive regulation of inflammatory response | 7.04E-03 | *Il1b/Il6/Ccl7/Ccl5/Setd4/Serpine1/Tnfsf18/Cd24* |
| GO:0048771 | Tissue remodeling | 9.08E-03 | *Ceacam1/Thbs4/Il6/Ccl2/Tnnt2/**Spp1**/Lif/Siglec15/Cd24* |

**Table S4. Distribution of fetal weights from SHRSP WT and SHRSP *Spp1*(Δ/Δ) litters at GD15.5 and 18.5.** Fetuses were stratified into percentile weight categories using the SHRSP WT fetal distribution as reference. Values are presented as number of fetuses and percentage of total fetuses per group. The proportion of growth restricted fetuses (< $10^{th}$ percentile) was compared between groups using Fisher's exact test.

| | GD 15.5 | | | GD 18.5 | | |
|---|---|---|---|---|---|---|
| | SHRSP WT | SHRSP *Spp1*(Δ/Δ) | *P* value | SHRSP WT | SHRSP *Spp1*(Δ/Δ) | *P* value |
| Number of Dams | 6 | 6 | | 3 | 6 | |
| Total Fetuses | 76 | 63 | | 28 | 62 | |
| < $10^{th}$ Percentile | 9.2% (7) | 11.1% (7) | 0.7813 | 7.1% (2) | 9.7% (6) | >0.9999 |
| 10-$90^{th}$ Percentile | 81.6% (62) | 81% (51) | | 85.8 (24) | 72.6% (45) | |
| > $90^{th}$ Percentile | 9.2% (7) | 7.9% (5) | >0.9999 | 7.1% (2) | 17.7% (11) | 0.2089 |

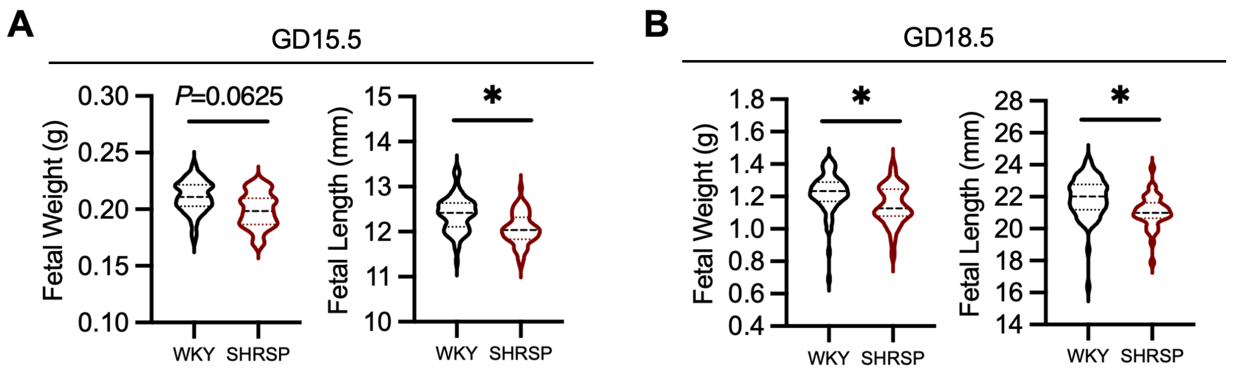


**Figure S1. Fetal weight and length in WKY and SHRSP pregnancies. (A)** Fetal weight and length at mid-gestation (GD15.5; WKY n=69 fetuses, N=6 dams; SHRSP n=57 fetuses, N=6 dams). **(B)** Fetal weight and length at late gestation (GD18.5; WKY n=72 fetuses, n=6 dams; SHRSP n=37 fetuses, n=4 dams). Asterisks denote statistical significance (*, $P<0.05$) using a linear mixed model to account for litter effects.

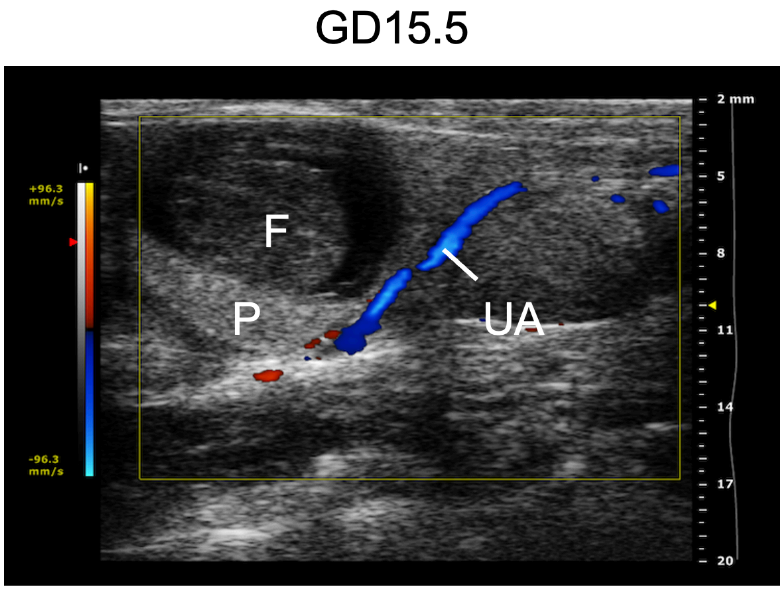


**Figure S2. Identification of the uterine artery at GD15.5 using Doppler ultrasound.** The uterine artery (UA) travels adjacent to the fetus (F) and placenta (P).

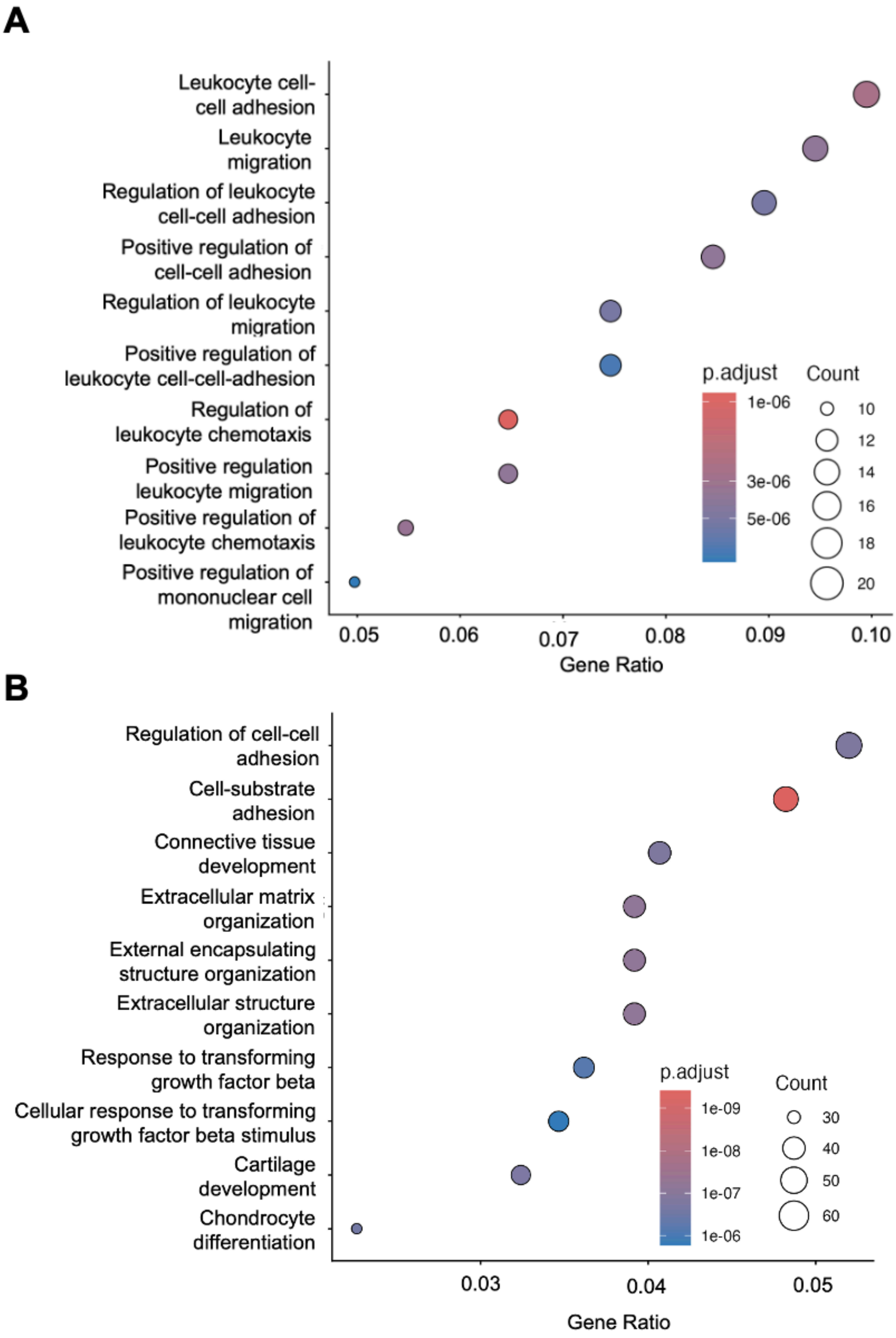
A
Leukocyte cell-cell adhesion
Leukocyte migration
Regulation of leukocyte cell-cell adhesion
Positive regulation of cell-cell adhesion
Regulation of leukocyte migration
Positive regulation of leukocyte cell-cell-adhesion
Regulation of leukocyte chemotaxis
Positive regulation leukocyte migration
Positive regulation of leukocyte chemotaxis
Positive regulation of mononuclear cell migration
0.05
0.06
0.07
0.08
0.09
0.10
Gene Ratio
p.adjust
1e-06
3e-06
5e-06
Count
10
12
14
16
18
20
B
Regulation of cell-cell adhesion
Cell-substrate adhesion
Connective tissue development
Extracellular matrix organization
External encapsulating structure organization
Extracellular structure organization
Response to transforming growth factor beta
Cellular response to transforming growth factor beta stimulus
Cartilage development
Chondrocyte differentiation
0.03
0.04
0.05
Gene Ratio
p.adjust
1e-09
1e-08
1e-07
1e-06
Count
30
40
50
60

**Figure S3. Additional biological process GO terms for SHRSP versus WKY uterine artery.** Bulk RNA sequencing was performed on GD15.5 WKY and SHRSP uterine arteries (N=5). **(A)** Top ten biological process GO terms for genes upregulated in SHRSP uterine arteries compared to WKY. **(B)** Top ten biological process GO terms for genes downregulated in SHRSP uterine arteries compared to WKY.

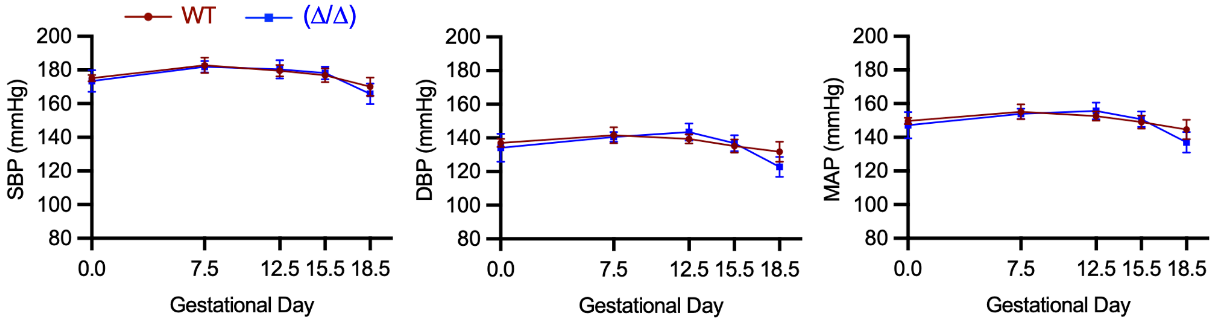


**Figure S4. OPN deficiency does not alter systemic blood pressure during hypertensive pregnancy.** Systolic, diastolic, and mean arterial pressure (SBP, DBP, MAP) across gestation in SHRSP WT and *Spp1*(Δ/Δ) pregnancy (N=5). Area under the curve analysis was used for statistics.

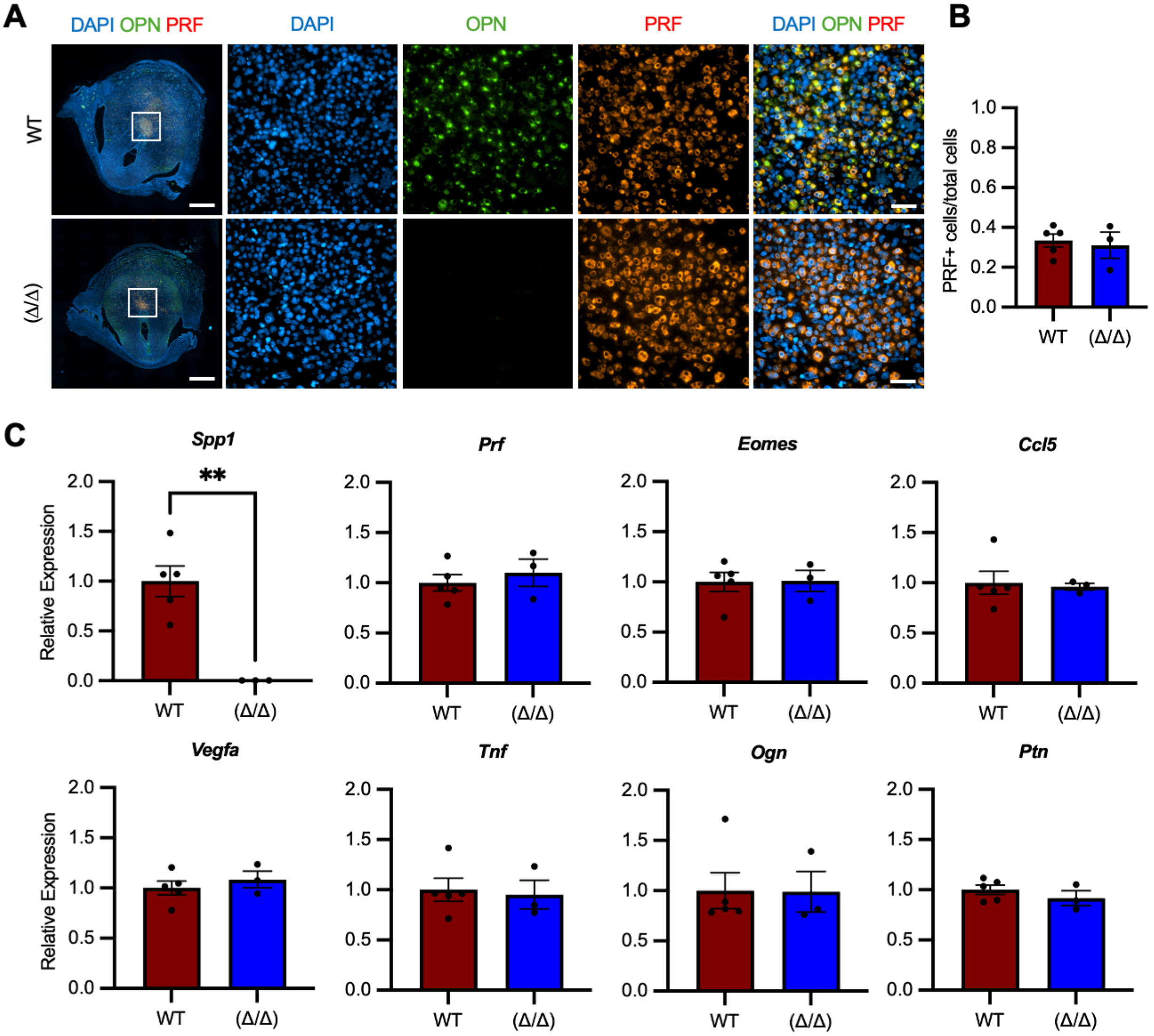


**Figure S5. OPN localizes to uterine NK cells, but OPN deficiency does not alter their abundance in the decidua at GD9.5. (A)** Low magnification (left panel) of whole SHRSP WT and *Spp1*(Δ/Δ) GD9.5 implantation sites showing osteopontin (OPN; green) and perforin (PRF; red). White box shows approximate location of high magnification images (right panels). Scale bar; 1000 μm (low magnification), 50 μm (high magnification). **(B)** Quantification of perforin-positive uterine NK cells in the ectoplacental cone region at GD9.5. **(C)** Relative expression of various genes associated with NK cell maturation in whole SHRSP WT and *Spp1*(Δ/Δ) GD9.5

implantation sites (N=3-5). Asterisks denote statistical significance (**, $P<0.01$) using Student's unpaired t-test.

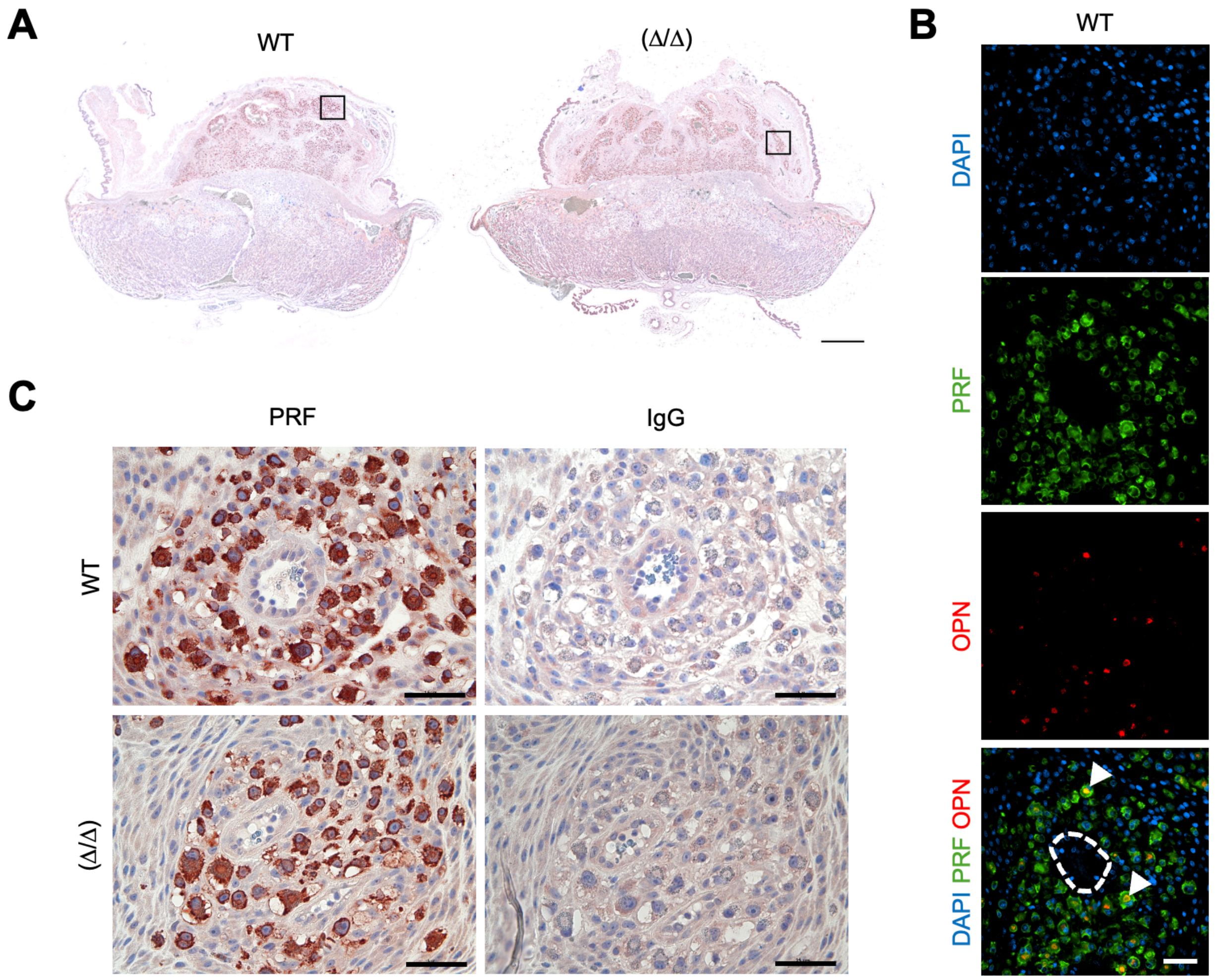


**Figure S6. OPN deficiency does not alter the abundance of uterine NK cells around spiral arteries at GD15.5. (A)** Low magnification GD15.5 whole placentas showing perforin (PRF)-positive uterine NK cells in the metrial gland using immunohistochemistry. Black box shows approximate location of (**B**) and (**C**). Scale bar; 1000 μm. **(B)** High magnification image showing osteopontin (OPN; red) co-localizing with PRF (green) in the metrial gland surrounding a spiral artery (outlined by dotted line) at GD15.5. Scale bar; 50 μm. **(C)** Immunohistochemistry showing PRF-positive uterine NK cells in metrial gland around spiral arteries in SHRSP WT and *Spp1*Δ/Δ. Scale bar; 50 μm.
